# Vertical and horizontal integration of multi-omics data with miodin

**DOI:** 10.1101/431429

**Authors:** Benjamin Ulfenborg

**Affiliations:** Department of Bioscience, University of Skövde, 544 28 Skövde, Sweden

**Keywords:** Multi-omics, data analysis, data integration, transparency

## Abstract

**Background:** Studies on multiple modalities of omics data such as transcriptomics, genomics and proteomics are growing in popularity, since they allow us to investigate complex mechanisms across molecular layers. It is widely recognized that integrative omics analysis holds the promise to unlock novel and actionable biological insights to health and disease. Integration of multi-omics data remains challenging, however, and requires combination of several software tools and extensive technical expertise to account for the properties of heterogeneous data.

**Results:** This paper presents the miodin R package, which provides a streamlined workflow-based syntax for multi-omics data analysis. The package allows users to perform analysis and integration of omics data either across experiments on the same samples, or across studies on the same variables. Workflows have been designed to promote transparent data analysis and reduce the technical expertise required to perform low-level data import and processing.

**Conclusions:** The miodin package is implemented in R and is freely available for use and extension under the GPL-3 license. Package source, reference documentation and user manual are available at https://gitlab.com/algoromics/miodin.

## Background

With the advances in high-throughput biotechnology over the past two decades, we now have access to an unprecedented wealth of data for many omics modalities. In this era of biomedical big data, the primary research challenges are how to integrate and analyze large-scale data of different types and sources to gain new insights into the complex mechanisms behind health and disease (1–4). In a study by Woo et al., DNA copy-number variation, methylation and gene expression were profiled in a cohort of hepatocellular carcinoma (HCC) patients. Integrative omics analysis revealed three molecular subtypes of HCC with differences in prognostic outcomes (5). Zhu et al. performed a comprehensive pan-cancer integrative analysis showing that a combination of clinical variables with molecular profiles improves prognostic power in 7 of the 14 cancer types studied (6). Lau et al. carried out a cardiac hypertrophy study in mice based on transcriptomics, proteomics and protein turnover data. Combining multi-omics data revealed complementary insights into the pathogenesis of the disease (7). These and other studies show that the integrative approach deliver novel biological insights. Advanced bioinformatics tools and algorithms have been developed that can analyze multiple modalities of omics data (8–10), but performing transparent and reproducible integrative analysis remains a significant challenge. Notably, considerable technical expertise is required to use many tools and combine them into a coherent pipeline.

Bioconductor is one of the largest open source projects for analysis of omics data (11), hosting more than 1 600 software packages as of release 3.8. Many experimental techniques (e.g. microarrays, sequencing and mass spectrometry) and omics data types (e.g. genomics, transcriptomics and proteomics) are supported (12–20). To perform data analysis the project hosts many packages for different workflow steps, such has import, annotation, preprocessing, quality control, statistical analysis, biological interpretation and visualization (12,21–26). By promoting a common set of data structures, package interoperability, version control, extensive documentation and high development standards, the project contributes significantly to distributing R software in bioinformatics. Furthermore, Bioconductor hosts experimental data, workflows, tutorials and other materials to facilitate learning, usage and combination of packages. With its large and active community, Bioconductor continues to expand to meet the future challenges in multi-omics data analysis.

Given the functionality it provides, Bioconductor is obvious choice when selecting software for performing integrative multi-omics data analysis. Though even for seasoned bioinformaticians, a lot of technical expertise and work is required to combine packages into coherent pipelines. Knowing which packages to use is an issue when working with new techniques and data, since there are several possible ones available for a given problem. Learning how to use several packages takes time, given the need to be familiar with the logic behind data structures along with multiple functions and their parameters. Another challenge is the growth in complexity of the analysis scripts, where code is required to perform every analysis step from import, through pre-processing, quality control, statistical analysis and interpretation. This problem is exacerbated when working with multi-omics data and performing integrated analysis, where several steps are needed for every experimental technique and data type. This increases the risk of clerical errors and results in low transparency in terms of what processing and analysis steps that have been performed.

A related problem in omics data analysis is the lack of a systematic way to specify generic study designs in analysis scripts. Issues may include what experimental variables to analyze, how to define sample groups and statistical comparisons, how samples are paired, any batch effects to consider and how to collapse replicates by mean. This can be performed ad hoc with e.g. variables and indexing operations but is error prone and gives low transparency when dealing with large datasets, multiple data types and more complex designs. Another general problem is the reproducibility of bioinformatics workflows (27,28), i.e. to ensure that the same results are obtained when running a workflow on the same data with the same steps and parameters. This has been addressed by Nextflow (29) and related software, which are used to construct workflows and support Docker technology (30) for deployment. This technology ensures that the analysis environment remains the same and protects against numeric instability across different systems. The BiocImageBuilder (31) is a tool designed to promote reproducibility of Bioconductor workflows by building a Docker image configured with all necessary software. The image also supports JupyterHub (32) and Binder (33) for private and public deployment of Jupyter notebooks for sharing and rerunning the analysis.

To address the challenges of multi-omics data analysis, the miodin R package was developed. The package provides a software infrastructure to build data analysis workflows that import, process and analyze multi-omics data. Workflows accommodate data of different omics modalities, including transcriptomics, genomics, epigenomics and proteomics, from different experimental techniques (microarrays, sequencing and mass spectrometry). The package allows users to integrate omics data from different experiments on the same samples (vertical integration) or across studies on the same variables (horizontal integration). Furthermore, the user is provided with an expressive vocabulary for declaring the experimental study design, to render this explicit within the analysis script and reduce the risk of clerical errors. A key design goal when developing miodin was to streamline data analysis, by providing a clean syntax for building workflows and minimizing the extent of technical expertise required to combine multiple software packages. The motivation behind this was to promote transparent biomedical data science.

## Implementation

### Package overview

The miodin package was implemented following the S4 object-oriented programming paradigm. Infrastructure functionality is supported by 16 S4 classes for which a common set of standard generics (base API) has been defined. Apart from the base classes, a number of workflow module classes have been developed, which serve as the building blocks of workflows. On top of the base API is a high-level user API consisting of convenience operators + and `%>%`along with helper functions to simply manipulation of objects (Figure 1). The user API has been developed to reduce the learning curve for the package and minimize the number of classes, functions and parameters the user needs to learn.

**Figure 1.**
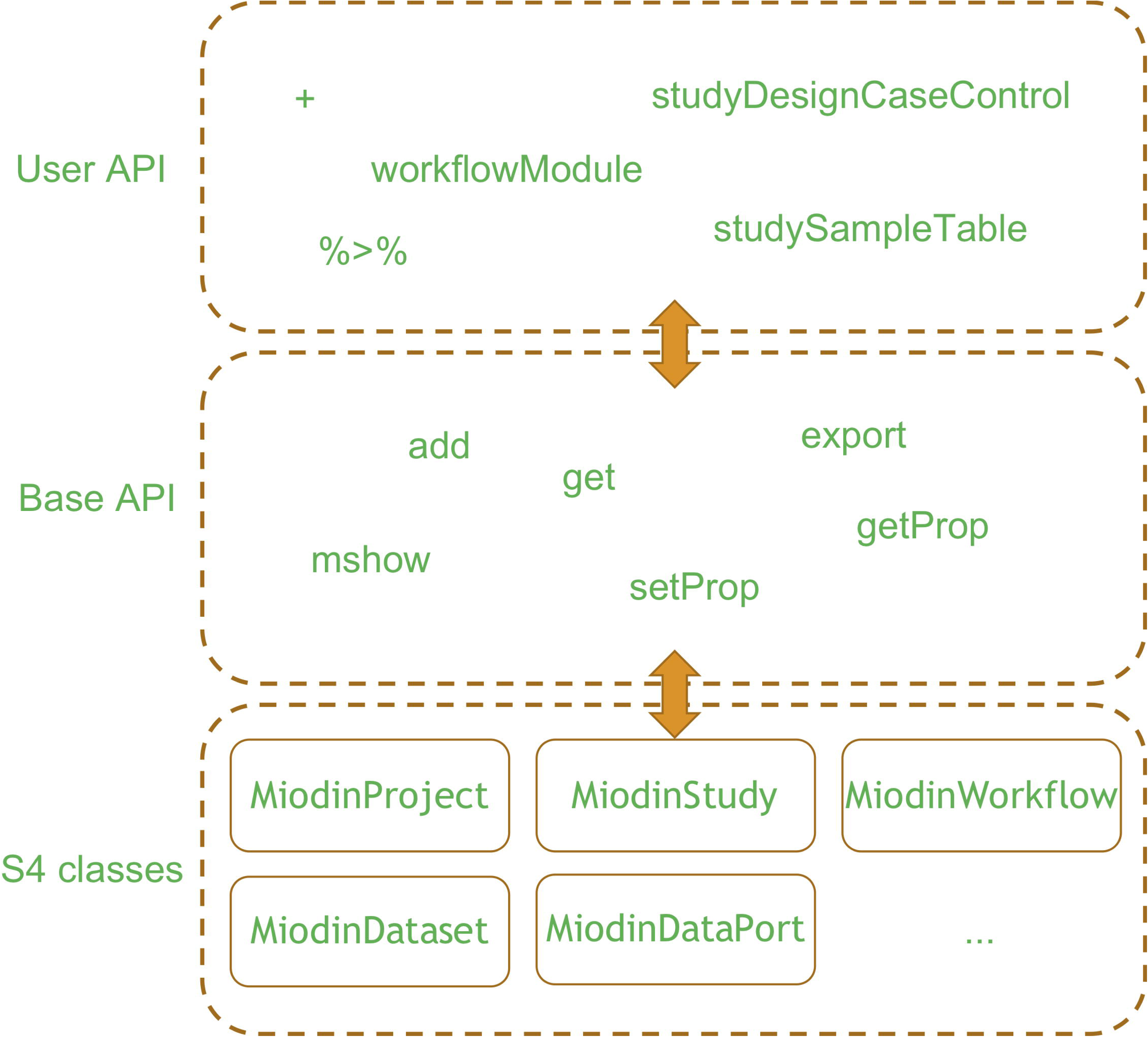
Package design. The miodin package provides a software infrastructure for data analysis implemented as a set of S4 classes. The base API contains standard generics for object manipulation and the user API provides convenience functions to facilitate package usage.

Data analysis with miodin follows an intuitive three-step process illustrated in Figure 2. The user first initializes a project, a study and a workflow. The project serves as a placeholder for all other objects and the study is used to declare the study design, including what samples and assays to analyze, and the experimental variables of interest, if any. The miodin package implements an expressive study design vocabulary and several convenience functions for common designs, such as case-control and time series experiments. These allow the user to declare all information required for data analysis in one place, thus reducing the risk of clerical errors in the analysis script and the amount of information the user must provide during the analysis itself. The workflow is used to build the data analysis procedure as a set of sequentially connected workflow modules that carry out specific tasks, such as data import or processing. The analysis is performed by executing the workflow, which generates datasets and results. These can be inspected, exported and used for further analysis.

**Figure 2.**
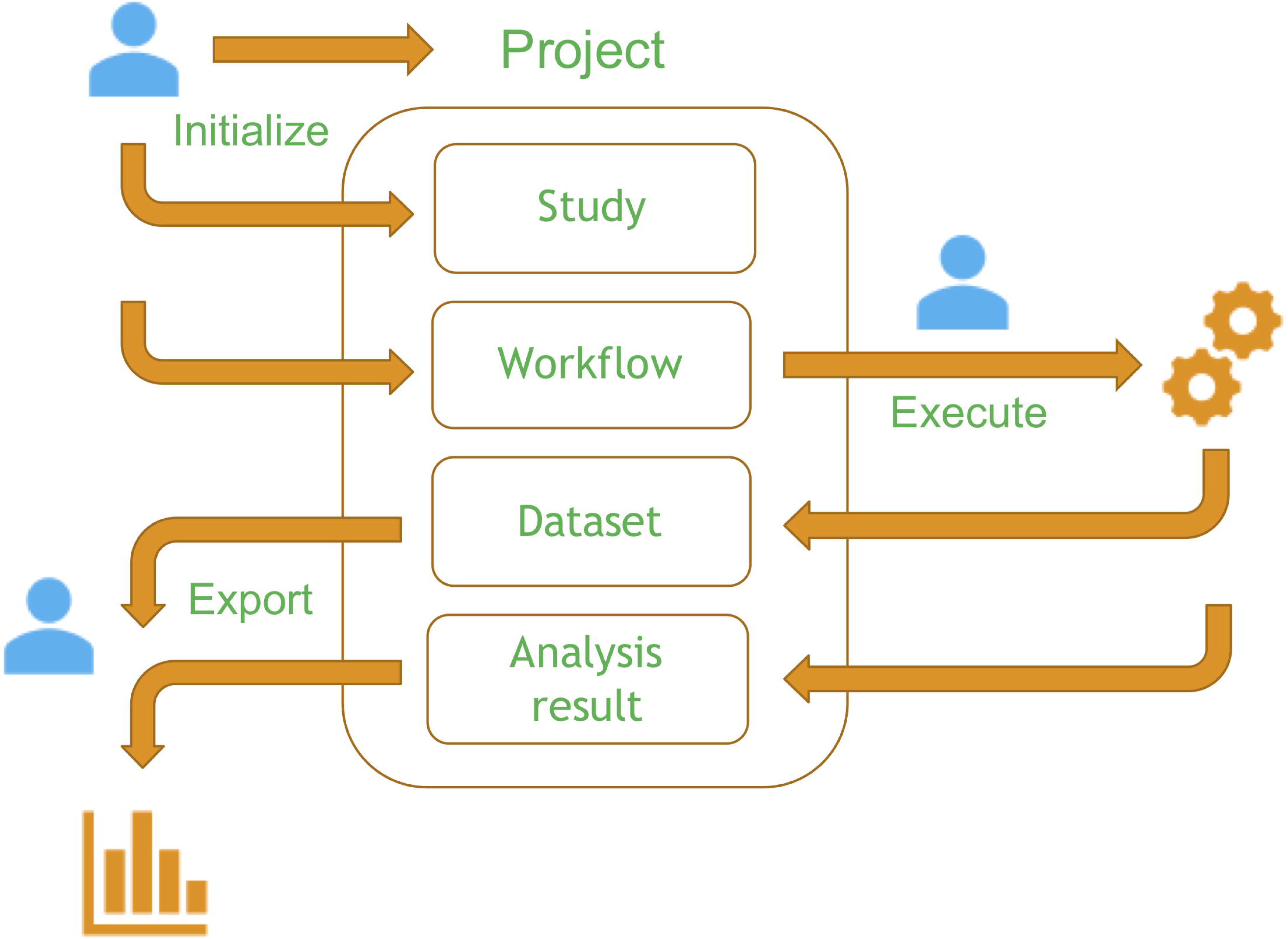
Data analysis in miodin. The user starts by defining a project, a study and a workflow. The study contains the design of the experiment and the workflow is defined by instantiating analysis modules, which generate datasets and analysis results upon execution. The user can then inspect and export the data and results.

### Study design vocabulary

Information related to a study is managed using the `MiodinStudy` class. The study design can be declared manually by instantiating an empty study and using helper functions that add different properties to the design (Table 1) or using convenience functions available for some of the most common designs (Table 2).

**Table 1:** Study design helper functions

| FUNCTION | DESCRIPTION |
| --- | --- |
| `studySamplingPoints` | Set the sampling points (e.g. time points) |
| `studyFactor` | Define a factor (experimental variable) |
| `studyGroup` | Define a sample group based on existing factors |
| `studyContrast` | Define a contrast (sample group comparison) |
| `studySampleTable` | Add a table with sample annotation data |
| `studyAssayTable` | Add a table with assay annotation data |

**Table 2:** Common study design functions

| FUNCTION | DESCRIPTION |
| --- | --- |
| `studyDesignCaseControl` | Single factor dividing samples into two groups |
| `studyDesignMultipleGroups` | Single factor dividing samples into multiple groups |
| `studyDesignRepeatedMeasures` | Single factor and multiple sampling points |
| `studyDesignTwoFactors` | Two factors and multiple sampling points |

The purpose of declaring the study design is for the user to give an explicit definition on what samples are included (in a sample table), what assays or experimental data files to analyze (in an assay table), what sample groups exist and which groups to compare during the analysis. The benefits of this are that the user can provide all this information in one place in the analysis script and that no variable manipulation is needed on the user’s part. Furthermore, when the user adds sample and assay tables these are automatically validated against the declared study design to detect potential clerical errors that might otherwise disturb the downstream analysis. For detailed examples how to declare the study design, see the miodin user manual in the GitLab repository (https://gitlab.com/algoromics/miodin).

### Workflow syntax

When the study design has been declared the next step is to define the data analysis workflow. A workflow is built by instantiating the `MiodinWorkflow` class and adding workflow modules to it, each one performing a specific task. Workflow modules are added to the workflow object by + operator and a module-specific instantiation function. To feed the output from one module as input to the next, they can be combined using the pipe operator `%>%`.

~~~
mw <-MiodinWorkflow(“DataAnalysisFlow”)
mw <-mw +
importMicroarrayData(…) %>%
processMicroarrayData(…) %>%
performOmicsModeling(…)
mw <-execute(mw)
~~~

This script initializes a workflow called DataAnalysisFlow with three workflow modules. Module parameters have been omitted for brevity. The first module imports microarray data, the second processes the output from the first module, and the final module performs statistical testing on the processed data. The analysis is carried out by calling the execute method. This syntax allows the user to define streamlined data analysis workflows, enhancing readability of the analysis script compared to longer chunks of code. Workflow modules have names starting with verbs denoting their function, making them easier to remember and improving analysis transparency. To further improve transparency, the analysis workflow automatically documents each processing and analysis step, including a description of what was done, what function was called, the name and version of the package, names and values of parameters, and how this affected the data. These can be inspected and exported as part of the dataset, thus addressing the issues of provenance (27), which is one important aspect of reproducibility.

### Package features

The workflow modules available as of miodin version 0.3.0 are described in Table 3. Import, processing and analysis of data is supported for different experimental techniques and omics data types as given in Table 4. For microarrays raw data from Affymetrix arrays (CEL format) and Illumina arrays (IDAT format) is supported, including transcriptomics, genomics (SNP) and methylation data. Processed data is also supported for microarrays, sequencing (RNA-seq counts) and mass spectrometry (protein quantification).

**Table 3:** Workflow modules

| FUNCTION | DESCRIPTION |
| --- | --- |
| `downloadRepositoryData` | Downloads data from an online repository |
| `importMicroarrayData` | Imports raw microarray data from Affymetrix and Illumina arrays |
| `importProcessedData` | Imports processed RNA, SNP, methylation and protein data |
| `processMicroarrayData` | Pre-processes microarray data |
| `processSequencingData` | Pre-processes sequencing data |
| `processMassSpecData` | Pre-processes mass spectrometry data |
| `integrateAssays` | Integrates several datasets into one |
| `performFactorAnalysis` | Performs factor analysis by matrix factorization |
| `performHypothesisTest` | Performs hypothesis testing |
| `performLinearModeling` | Performs linear or generalized linear modeling |
| `performOmicsModeling` | Performs modeling with limma, snpStats, DMRcate or edgeR depending on the input object |

**Table 4:** Supported experimental techniques and data types

|  | RNA | SNP | Methylation | Protein |
| --- | --- | --- | --- | --- |
| Microarray | Raw and processed | Raw and processed | Raw and processed |  |
| Sequencing | Processed |  |  |  |
| Mass spectrometry |  |  |  | Processed |

### Omics data integration

The miodin package can be used for analysis single-omics data, though by design the package is intended to streamline multi-omics data integration and analysis. Two case studies were carried out to illustrate how horizontal integration (across studies) and vertical integration (across omics modalities) can be performed with miodin. Three lung cancer transcriptomics datasets with accession number E-GEOD-27262 (34), E-GEOD-19188 (35) and E-GEOD-40791 (36) were downloaded from ArrrayExpress (37) and prepared for horizontal integration. Probes were mapped to genes with NetAffx file HG-U133-Plus-2-NA36 and each dataset was filtered to include only the first 2000 genes. Genes with an expression below 5 in all samples were removed. Differentially expressed genes between tumor samples and non-tumorigenic controls were identified with limma (19), using adjusted p-value < 0.05 and absolute log2 fold change > 1. The analysis script is available in Additional file 1.

Vertical integration was carried out using breast cancer data from the curatedTCGAData package (38). RNA-seq gene and miRNA count data as well as 450k methylation data were integrated and used to fit a Multi-Omics Factor Analysis (MOFA) model (39). RNA-seq gene and methylation data were filtered to include the 5000 top-variance features. Both RNA-seq datasets were subject to variance stabilization normalization with DESeq2 (40). Methylation probes flagged as problematic by DMRcate (41) and non-CpG probes were removed. The analysis script is supplied in Additional file 2. Jupyter notebooks for reproducing the analysis are provided in GitLab (https://gitlab.com/algoromics/miodin-notebooks), with the option to run on Binder (33).

## Results

### Horizontal integration: meta-analysis on lung cancer transcriptomics data

To perform meta-analysis in miodin, a study design must be declared for every dataset included in the analysis. This implies defining a sample table and assay table (as data frames) and calling the appropriate study design function. The three transcriptomics datasets used here (referred to as Wei2012, Hou2010 and Zhang2012) have case-control designs and were declared using `studyDesignCaseControl` (see Additional file 1). The Wei2012 dataset contained 50 samples; 25 from stage 1 lung adenocarcinoma tissue and 25 paired samples from adjacent normal tissue (34). Sample pairedness was specified with the `paired` argument to the study design function, naming a column in the sample table containing information of how samples are paired. The Hou2010 dataset contained 156 samples (91 tumors and 65 healthy controls) and Zhang2012 contained 194 samples (94 tumors and 100 healthy controls).

When the study design had been declared, a workflow was built to import and process transcriptomics data. The raw microarray data from the three datasets has been normalized, annotated and included as part of the companion package miodindata, so this workflow imported data using `importProcessedData` and performed further processing with `processMicroarrayData`. The three datasets were integrated using `integrateAssays` and linear modeling with limma was carried out with `performOmicsModeling`. This module identifies differentially expressed genes in each individual dataset and by setting `metaAnalysis` to `TRUE` an additional step is performed to reveal concordant results between the datasets. The final results are stored as a Venn diagram accompanied by data frames, which can be exported to disk (data frames are exported as Excel sheets). The Venn diagram is shown in Figure 3 and the list of differentially expressed genes is provided in Additional file 3.

**Figure 3.**
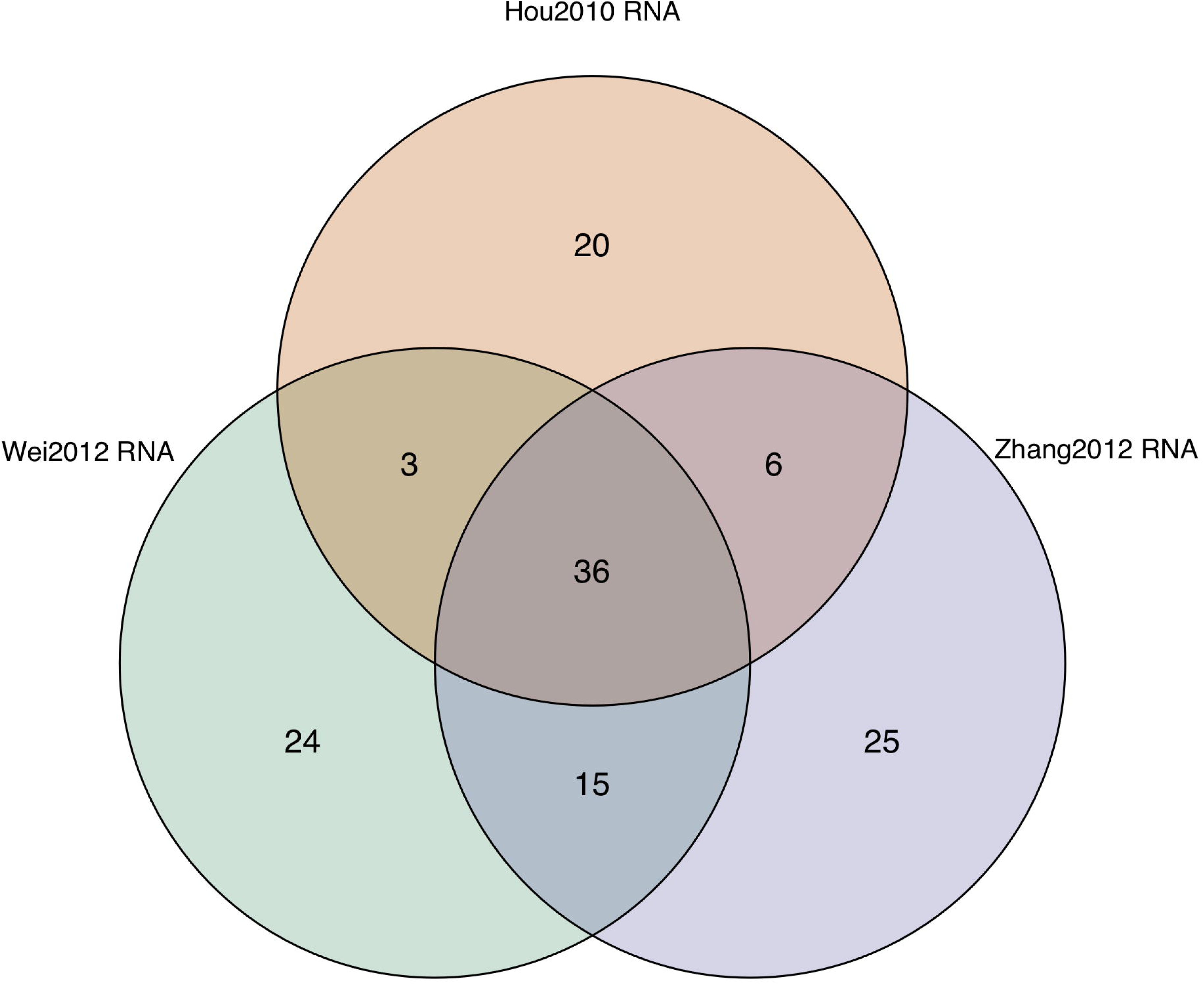
Venn diagram of the number of genes identified as differentially expressed in the Wei2012, Hou2010 and Zhang 2012 datasets.

### Vertical integration: exploratory data analysis on multi-omics data

The TCGA data used for vertical integration contained 338 breast cancer samples, for which survival status (alive or diseased) was available. RNA-seq gene and miRNA as well as 450k methylation data was processed and included in the miodindata package to ease replication of the analysis workflow. To perform vertical integration, a case-control study design was declared and one assay table for each omics modality was added, specifying the data files to import. A workflow was built to import data with `importProcessedData`, followed `processSequencingData` to perform RNA-seq count filtering and variance stabilization. Methylation data was processed with `processMicroarrayData` to remove bad and non-CpG probes. The multi-omics data was integrated with `integrateAssays` and integrative analysis carried out with `performFactorAnalysis` (see Additional file 2). This module runs MOFA, which is a matrix factorization technique for inferring latent factors that account for omics-specific and shared variance across omics modalities. Two strengths of MOFA are that it can integrate data from different distributions and handle missing data (39).

The results from `performFactorAnalysis` include the fitted model object and plots to assess the model in terms of variance explained, sample clustering (Figure 4) and the top features in the first factor (Figure 5). Plots for other factors can be rendered and further downstream analysis carried out with the model object.

**Figure 4.**
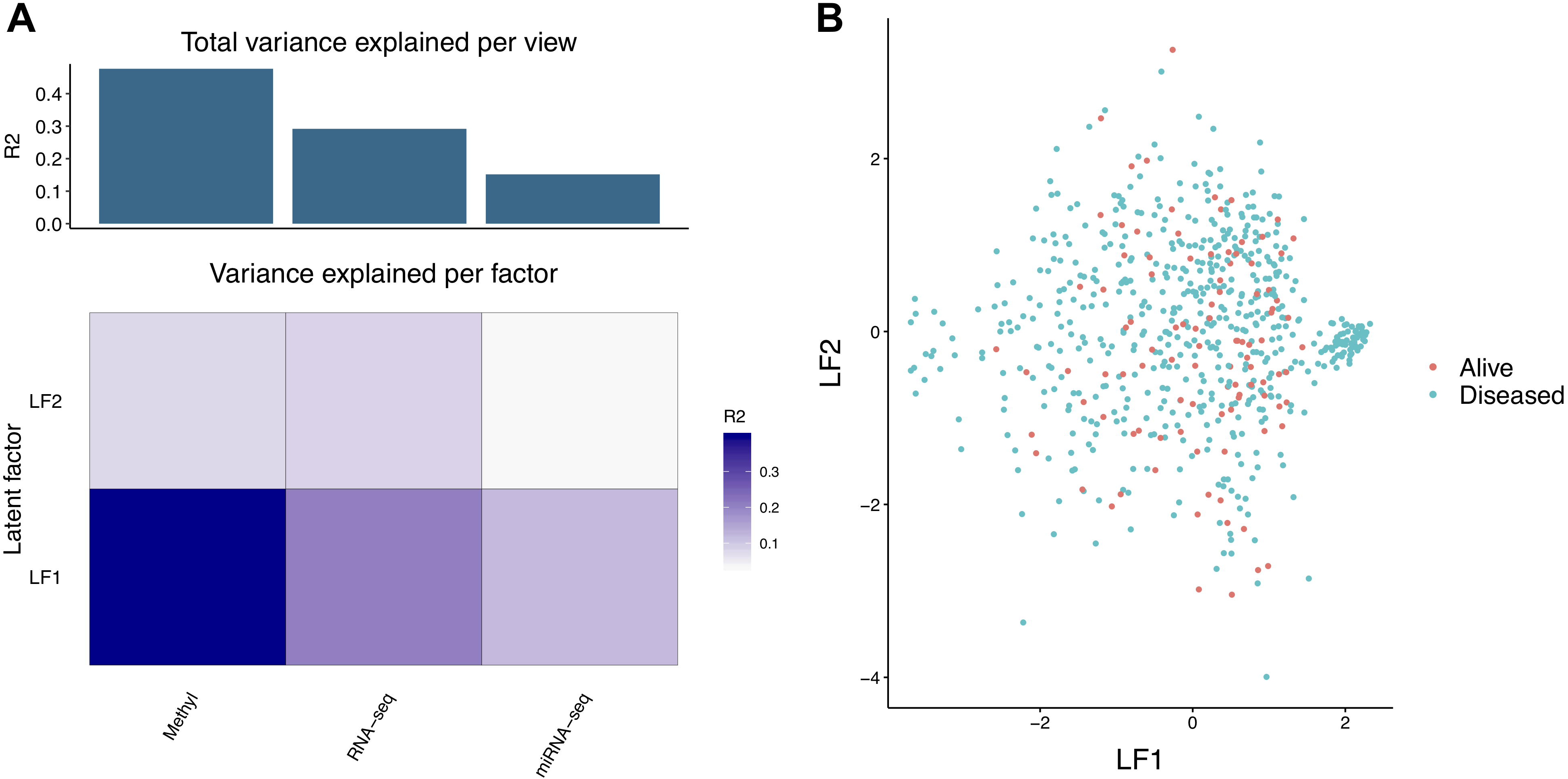
Assessment of the fitted MOFA model. Panel A shows the total amount of variance explained by the model in each omics modality (view) and variance explained per factor. Panel B shows a sample ordination plot based on latent factor (LF) 1 and 2.

**Figure 5.**
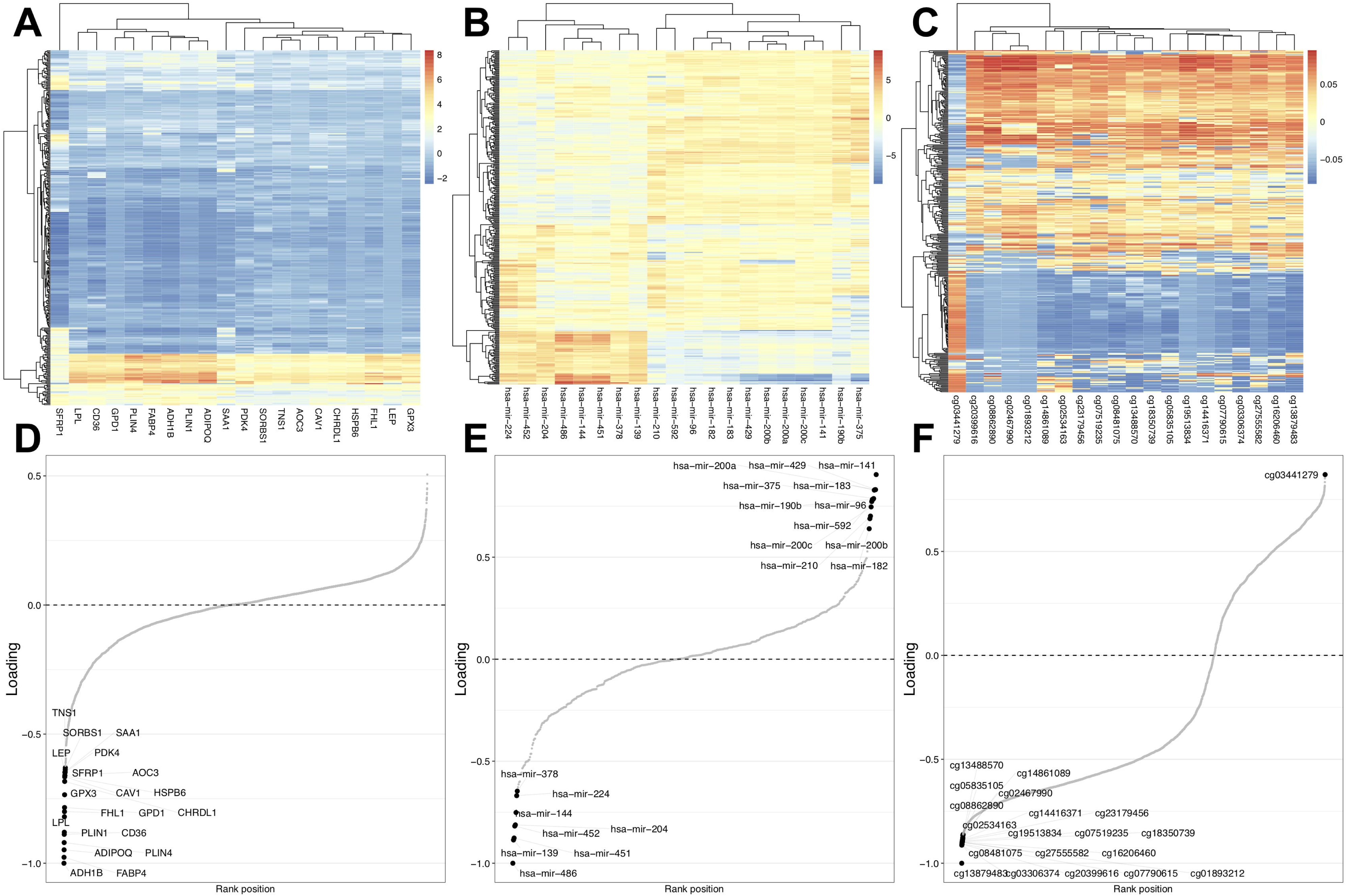
Interpretation of MOFA factor 1. Panels A through C show sample heatmaps with the top features in the factor for RNA-seq gene, miRNA and 450k methylation data, respectively. Panels D through F reveal the loadings of the top features corresponding to the heatmaps in Panels A through C.

## Discussion

Multi-omics experiments are indispensable for studying biological systems across molecular layers and tools for performing data analysis are actively being developed (42–45). One outstanding challenge in multi-omics data integration and analysis is to combine several tools into a coherent pipeline, which is time-consuming and requires extensive technical knowledge of various data types and platforms. The miodin package implements an infrastructure for addressing this challenge, by providing a high-level user API for declaring study designs and building analysis workflows with support for multiple omics data types and platforms. A key design goals was to streamline data analysis and lower the technical expertise and time required to perform low-level processing steps. This way users can spend more time on performing the actual analysis and interpreting results. The package comes with extensive documentation and examples, to render it readily accessible and ensure it benefits the multiomics research community.

Steps have also been taken to promote transparent and reproducible data analysis. Apart from streamlined workflow syntax, processing and analysis steps are automatically documented when workflows are executed, and a Docker image has been prepared for containerized deployment of Jupyter notebooks. Other tools such as Nextflow (29) have previously addressed the issues of building computational workflows and ensuring reproducibility. Nextflow is extremely versatile as it can combine command line tools and scripts in different programming languages into pipelines that are executed in containers. Furthermore, support is provided for deployment of workflows on computer clusters and cloud computing services, making workflow execution parallelized and scalable. The miodin package, on the other hand, provides an infrastructure for multi-omics data that streamlines import, processing, integration and analysis. The workflows developed with miodin can readily be incorporated into Nextflow pipelines to combine them with other computational tools, and to leverage Nextflow’s massive parallelization and scalability.

Several future developments are planned to enhance the functionality of the miodin package. Firstly, additional omics data types and platforms (e.g. raw sequencing and proteomics data, metabolomics, single cell, qPCR) will be supported. Secondly, several statistical and high-level analysis methods (e.g. clustering, classification, networks, annotation enrichment) will be implemented. Thirdly, workflow modules will be added for obtaining data from additional public repositories for omics, interaction and annotation data. Finally, support for automated deployment of workflows on remote computer resources will be implemented, to leverage the parallelization and scalability of computer clusters and cloud services.

## Conclusions

This paper presented the miodin package, which provides an infrastructure for integration and analysis of multi-omics data. Key features include a high-level user API, an expressive vocabulary for declaring study designs, streamlined workflows and support for multiple omics data types and platforms. The package has been designed to promote transparent data analysis and a Docker image has been prepared for deployment of Jupyter notebooks to ensure workflow reproducibility. Notebooks can also be executed on Binder, which provides an accessible web-based interface for developing and testing workflows. To ensure the research community benefits from miodin, the software package with extensive documentation is made freely available on GitLab under the GPL-3 license.

### Availability and requirements

**Project name:** miodin

**Project home page:** https://gitlab.com/algoromics/miodin

**Operating system(s):** Windows, Linux, MacOS

**Programming language:** R

**Other requirements:** Python

**License:** GNU General Public License v3.0

**Any restrictions to use by non-academics:** No

### Additional files

Additional file 1: **Horizontal integration analysis script.** R script for performing horizontal integration as presented in the paper. (R 5 kb)

Additional file 2: **Vertical integration analysis script.** R script for performing vertical integration as presented in the paper. (R 3 kb)

Additional file 3: **Differentially expressed genes from meta-analysis.** List of genes found differentially expressed in horizontal integration analysis. (XLSX 18 kb)

#### List of abbreviations

HCC: hepatocellular carcinoma
MOFA: Multi-Omics Factor Analysis
TCGA: The Cancer Genome Atlas
LF: Latent factor

## Declarations

### Ethics approval and consent to participate

Not applicable.

### Consent for publication

Not applicable.

### Availability of data and material

Source code and user manual for the miodin package are available on GitLab (https://gitlab.com/algoromics/miodin). Additional file 1 contains the analysis script for horizontal integration. Additional file 2 contains the analysis script for vertical integration. Additional file 3 contains the list of differentially expressed genes identified in horizontal integration analysis. Processed datasets used for analysis are available as part of the miodindata companion package, also available on GitLab (https://gitlab.com/algoromics/miodindata). Source datasets for horizontal integration are available from ArrayExpress with accession numbers E-GEOD-27262, E-GEOD-19188 and E-GEOD-40791. Source datasets for vertical integration are available in the curatedTCGAData package from Bioconductor, <u>10.18129/B9.bioc.curatedTCGAData</u>.

### Authors’ contributions

Not applicable.

## Acknowledgements

Preparation of data for vertical integration was performed on resources provided by the Swedish National Infrastructure for Computing (SNIC) at Uppsala Multidisciplinary Center for Advanced Computational Science (UPPMAX).

## References

1. Joyce AR, Palsson BØ. The model organism as a system: integrating “omics” data sets. Nat Rev Mol Cell Biol. 2006;7(3):198–210.

2. Ebrahim A, Brunk E, Tan J, O’Brien EJ, Kim D, Szubin R, et al. Multi-omic data integration enables discovery of hidden biological regularities. Nat Commun. 2016;7:1–9.

3. Berger B, Peng J, Singh M. Computational solutions for omics data. Nat Rev Genet. 2013;8(9):1385–95.

4. Karczewski KJ, Snyder MP. Integrative omics for health and disease. Nat Rev Genet. 2018;19(5):29–39.

5. Woo HG, Choi JH, Yoon S, Jee BA, Cho EJ, Lee JH, et al. Integrative analysis of genomic and epigenomic regulation of the transcriptome in liver cancer. Nat Commun. 2017;8(1).

6. Zhu B, Song N, Shen R, Arora A, Machiela MJ, Song L, et al. Integrating Clinical and Multiple Omics Data for Prognostic Assessment across Human Cancers. Sci Rep. 2017;7(1):1–13.

7. Lau E, Cao Q, Lam MPY, Wang J, Ng DCM, Bleakley BJ, et al. Integrated omics dissection of proteome dynamics during cardiac remodeling. Nat Commun. 2018;9(1).

8. Reich M, Liefeld T, Gould J, Lerner J, Tamayo P, Mesirov JP. GenePattern 2.0. Nat Genet. 2006;38(5).

9. Fisch KM, Meißner T, Gioia L, Ducom JC, Carland TM, Loguercio S, et al. Omics Pipe: A community-based framework for reproducible multi-omics data analysis. Bioinformatics. 2015;31(11):1724–8.

10. Afgan E, Baker D, Batut B, Van Den Beek M, Bouvier D, Čech M, et al. The Galaxy platform for accessible, reproducible and collaborative biomedical analyses: 2018 update. Nucleic Acids Res. 2018;46(W1):W537–W544.

11. Huber W, Carey VJ, Gentleman R, Anders S, Carlson M, Carvalho BS, et al. Orchestrating high-throughput genomic analysis with Bioconductor. Nat Methods. 2015;12(2):115–21.

12. Carvalho BS, Irizarry R a. A framework for oligonucleotide microarray preprocessing. Bioinformatics. 2010;26(19):2363–7.

13. Aryee MJ, Jaffe AE, Corrada-Bravo H, Ladd-Acosta C, Feinberg AP, Hansen KD, et al. Minfi: a flexible and comprehensive Bioconductor package for the analysis of Infinium DNA methylation microarrays. Bioinformatics. 2014 May 15;30(10):1363–9.

14. Dvinge H, Bertone P. HTqPCR: high-throughput analysis and visualization of quantitative real-time PCR data in R. Bioinformatics. 2009 Dec 15;25(24):3325–6.

15. Gatto L, Lilley KS. MSnbase-an R/Bioconductor package for isobaric tagged mass spectrometry data visualization, processing and quantitation. Bioinformatics. 2012;28(2):288–9.

16. Hahne F, LeMeur N, Brinkman RR, Ellis B, Haaland P, Sarkar D, et al. flowCore: a Bioconductor package for high throughput flow cytometry. BMC Bioinformatics. 2009 Dec 9;10(1):106.

17. Lawrence M, Gentleman R. VariantTools: an extensible framework for developing and testing variant callers. Bioinformatics. 2017;33(20):3311–3.

18. Liao Y, Smyth GK, Shi W. The Subread aligner: fast, accurate and scalable read mapping by seed-and-vote. Nucleic Acids Res. 2013;41(10):e108–e108.

19. Ritchie ME, Phipson B, Wu D, Hu Y, Law CW, Shi W, et al. limma powers differential expression analyses for RNA-sequencing and microarray studies. Nucleic Acids Res. 2015;1–13.

20. Wehrens R, Weingart G, Mattivi F. metaMS: An open-source pipeline for GC--MS-based untargeted metabolomics. J Chromatogr B. 2014;966:109–16.

21. Gentleman R. Annotate: Annotation for microarrays. R package version 1.56. 1. 2016.

22. Leek JT, Johnson WE, Parker HS, Jaffe AE, Storey JD. The sva package for removing batch effects and other unwanted variation in high-throughput experiments. Bioinformatics. 2012;28(6):882–3.

23. Morgan M, Anders S, Lawrence M, Aboyoun P, Pages H, Gentleman R. ShortRead: a bioconductor package for input, quality assessment and exploration of high-throughput sequence data. Bioinformatics. 2009;25(19):2607–8.

24. Robinson MD, McCarthy DJ, Smyth GK. edgeR: a Bioconductor package for differential expression analysis of digital gene expression data. Bioinformatics. 2010;26(1):139–40.

25. Falcon S, Gentleman R. Using GOstats to test gene lists for GO term association. Bioinformatics. 2007 Jan 15;23(2):257–8.

26. Luo W, Brouwer C. Pathview: an R/Bioconductor package for pathway-based data integration and visualization. Bioinformatics. 2013 Jul 15;29(14):1830–1.

27. Kanwal S, Khan FZ, Lonie A, Sinnott RO. Investigating reproducibility and tracking provenance - A genomic workflow case study. BMC Bioinformatics. 2017;18(1):1–14.

28. Kulkarni N, Alessandrì L, Panero R, Arigoni M, Olivero M, Ferrero G, et al. Reproducible bioinformatics project: A community for reproducible bioinformatics analysis pipelines. BMC Bioinformatics. 2018;19(Suppl 10).

29. Di Tommaso P, Chatzou M, Floden EW, Barja PP, Palumbo E, Notredame C. Nextflow enables reproducible computational workflows. Nat Biotechnol. 2017;35(4):316–9.

30. Merkel D. Docker: lightweight linux containers for consistent development and deployment. Linux J. 2014;2014(239):2.

31. Almugbel R, Hung LH, Hu J, Almutairy A, Ortogero N, Tamta Y, et al. Reproducible Bioconductor workflows using browser-based interactive notebooks and containers. J Am Med Informatics Assoc. 2018;25(1):4–12.

32. Ragan-Kelley M, Kelley K, Kluyver T. JupyterHub: deploying Jupyter notebooks for students and researchers. 2019.

33. Binder [Internet]. 2019 [cited 2019 Feb 2]. Available from: https://mybinder.org

34. Wei TYW, Juan CC, Hisa JY, Su LJ, Lee YCG, Chou HY, et al. Protein arginine methyltransferase 5 is a potential oncoprotein that upregulates G1 cyclins/cyclin-dependent kinases and the phosphoinositide 3-kinase/AKT signaling cascade. Cancer Sci. 2012;103(9):1640–50.

35. Hou J, Aerts J, den Hamer B, van Ijcken W, den Bakker M, Riegman P, et al. Gene expression-based classification of non-small cell lung carcinomas and survival prediction. PLoS One. 2010;5(4):e10312.

36. Zhang Y, Foreman O, Wigle DA, Kosari F, Vasmatzis G, Salisbury JL, et al. USP44 regulates centrosome positioning to prevent aneuploidy and suppress tumorigenesis. J Clin Invest. 2012;122(12):4362–74.

37. Parkinson H, Kapushesky M, Shojatalab M, Abeygunawardena N, Coulson R, Farne A, et al. ArrayExpress - A public database of microarray experiments and gene expression profiles. Nucleic Acids Res. 2007;35(Database issue):D747–50.

38. Ramos M, Waldron L, Schiffer L, Obenchain V, Martin M. curatedTCGAData: Curated Data From The Cancer Genome Atlas (TCGA) as MultiAssayExperiment Objects. R Packag version 120. 2018;

39. Argelaguet R, Velten B, Arnol D, Dietrich S, Zenz T, Marioni JC, et al. Multi-Omics Factor Analysis—a framework for unsupervised integration of multi-omics data sets. Mol Syst Biol. 2018;v14(e8124):1–13.

40. Love MI, Huber W, Anders S. Moderated estimation of fold change and dispersion for RNA-seq data with DESeq2. Genome Biol. 2014;15(12):1–21.

41. Peters TJ, Buckley MJ, Statham AL, Pidsley R, Samaras K, V Lord R, et al. De novo identification of differentially methylated regions in the human genome. Epigenetics and Chromatin. 2015;8(1):1–16.

42. Vantaku V, Dong J, Ambati CR, Perera D, Donepudi SR, Amara CS, et al. Multi-omics integration analysis robustly predicts high-grade patient survival and identifies CPT1B effect on fatty acid metabolism in Bladder Cancer. Clin Cancer Res. 2019.

43. Huang L, Brunell D, Stephan C, Mancuso J, Yu X, He B, et al. Driver Network as a Biomarker?: Systematic integration and network modeling of multi-omics data to derive driver signaling pathways for drug combination prediction. Bioinformatics. 2019.

44. Dao MC, Sokolovska N, Brazeilles R, Affeldt S, Pelloux V, Prifti E, et al. A Data Integration Multi-Omics Approach to Study Calorie Restriction-Induced Changes in Insulin Sensitivity. Front Physiol. 2019;9(February).

45. Chung NC, Mirza B, Choi H, Wang J, Wang D, Ping P, et al. Unsupervised Classification of Multi-Omics Data during Cardiac Remodeling using Deep Learning. Methods. 2019.

